# The impact of genetic risk for Alzheimer’s disease on the structural brain networks of young adults

**DOI:** 10.1101/2021.09.22.461338

**Authors:** Anastasia Mirza-Davies, Sonya Foley, Xavier Caseras, Emily Baker, Peter Holmans, Valentina Escott-Price, Derek K. Jones, Judith R. Harrison, Eirini Messaritaki

## Abstract

We investigated the structural brain networks of 562 young adults in relation to polygenic risk for Alzheimer’s disease, using magnetic resonance imaging (MRI) and genotype data from the Avon Longitudinal Study of Parents and Children. Diffusion MRI data were used to perform whole-brain tractography and to generate structural brain networks for the whole-brain connectome, and for the default mode, limbic and visual subnetworks. The mean clustering coefficient, mean betweenness centrality, characteristic path length, global efficiency and mean nodal strength were calculated for these networks, for each participant. The connectivity of the rich-club, feeder and local connections was also calculated. Polygenic risk scores (PRS), estimating each participant’s genetic risk, were calculated at genome-wide level and for nine specific disease pathways. Correlations were calculated between the PRS and a) the graph theoretical metrics of the structural networks and b) the rich-club, feeder and local connectivity of the whole-brain networks.

In the visual subnetwork, the mean nodal strength was negatively correlated with the genomewide PRS (r=−0.19, *p*=1.3×10^−5^), the mean betweenness centrality was positively correlated with the plasma lipoprotein particle assembly PRS (r=0.16, *p*=9.2×10^−4^), and the mean clustering coefficient was negatively correlated with the tau protein binding PRS (r=−0.16, *p*=9.2×10^−4^). In the default mode network, the mean nodal strength was negatively correlated with the genomewide PRS (r=−0.14, *p*=1.5×10^−3^). The rich-club and feeder connectivities were negatively correlated with the genome-wide PRS (r=−0.16, *p*=3.7×10^−4^; r=−0.15, *p*=8.8×10^−4^). Our results indicate small reductions in brain connectivity in young adults at risk of developing Alzheimer’s disease in later life.

## 1. Introduction

Alzheimer’s disease (AD) is a progressive neurodegenerative disorder that affects over 35 million people world-wide (Prince et al., 2013). It leads to severe cognitive impairment and the inability of patients to function independently. There is a pressing need to identify non-invasive biomarkers that could facilitate pre-symptomatic diagnosis when disease-modifying therapies become available. Although a minority of early-onset AD cases are caused by mutations in specific genes with autosomal dominant inheritance (Tanzi, 2012), the majority of AD has a complex genetic architecture and is highly heritable (Gatz et al., 2006), with different genes conveying different amounts of risk. Genome-wide Association Studies (GWAS) have implicated many Single Nucleotide Polymorphisms (SNPs) (Kunkle et al., 2019), of which the apolipoprotein e4 allele *(APOE4)* confers the greatest risk (Strittmatter et al., 1993; Saunders et al., 1993; Lambert et al., 2013; Yu et al., 2014; Farrer et al., 1997), but is neither necessary nor sufficient to cause AD (Sims et al, 2020). AD GWAS have also found evidence that specific biological processes, or disease pathways, such as cell trafficking, beta amyloid production, tau protein regulation and cholesterol transport are involved (Kunkle et al., 2019; Jones et al., 2010). Polygenic risk scores (PRS), which aggregate risk loci genome-wide (Wray et al., 2014), are highly predictive of AD (Sleegers et al., 2015; Xiao et al., 2015; Yokohama et al., 2015; Escott-Price et al., 2015; Escott-Price et al., 2017; Tosto et al., 2017; Chaudhury et al., 2018; Cruchaga et al., 2018; Harrison et al., 2020; Altmann et al., 2020) and have been widely used in the search for biomarkers for the disease (Harrison et al., 2020).

Obtaining reliable biomarkers in a non-invasive manner is very valuable because it can be better tolerated by participants compared to more invasive methods (Prestia et al., 2013; Zhang et al., 2012). Magnetic resonance imaging (MRI) can non-invasively measure characteristics of the brain’s structure. Diffusion-weighted MRI (dMRI, Le Bihan et al., 2006) has allowed mapping of the brain’s white-matter (WM) tracts, enabling the study of the human brain as a network of cortical and subcortical areas connected via those tracts. Via these techniques, alterations in the brain of AD patients and of people at risk of developing AD have been identified. AD patients exhibit axonal loss in tracts associated with certain default mode network (DMN) nodes (Mito et al., 2018). They also exhibit increased characteristic path length and decreased intramodular connections in functional and structural brain networks compared to healthy controls (Dai et al., 2018). The DMN is altered in the presence of AD pathology (Dai et al., 2018) where a decrease in its connectivity has been observed (Mohan et al., 2016; Badhwar et al., 2017). The diffusion tensor fractional anisotropy in the cingulum and of the splenium of the corpus callosum is reduced in AD patients compared to controls (Zhang et al., 2007). Structural covariance brain networks, in which the edges are calculated as the correlations between the node volumes, show decreased small-worldness in AD (John et al, 2017). Increased shortest path length and clustering coefficient, as well as decreased global and local efficiency have been observed in the structural brain networks of AD patients (He et al., 2008; Lo et al., 2010). These results, as well as recent work by Palesi et al. (2016), suggest that, in addition to the AD pathology preferentially affecting specific brain areas, AD is a disconnection syndrome.

Cognitively healthy middle-aged and older carriers of AD risk (genetic or otherwise) also exhibit alterations in brain structure. Decreased hippocampal volume and cortical thickness have been associated with high AD PRS (Mormino et al., 2016; Corlier et al., 2017; Li et al., 2017). Ageing *APOE4* carriers have reduced local structural connectivity at the precuneus, medial orbitofrontal cortex and lateral parietal cortex (Brown et al., 2011). *APOE4* status also affects the clustering coefficient and the local efficiency of structural brain networks. Specifically, Ma et al. (2017) observed that the values for the *APOE4* carriers were higher than those of the non-carriers in a normal-cognition group, while the opposite pattern was observed in a group of participants suffering from Mild Cognitive Impairment (MCI). Middle-aged adults with genetic, family and lifestyle risks of developing AD have a hub in their structural connectome that is not present in the structural connectome of people with no such risks of developing AD (Clarke et al., 2020). Significant functional connectivity differences in the brain networks implicated in cognition were seen in middle-aged individuals with a genetic risk for AD (Goveas et al., 2013). The DMN also exhibits changes in mature (Fleisher et al., 2009) and young *APOE4* carriers (Filippini et al., 2009). A PRS composed of immune risk SNPs is associated with a thinner regional cortex in healthy older adults at risk of developing AD (Corlier et al., 2017). Other studies have also investigated the effect of AD PRS on brain structure (Lupton et al., 2016; Hayes et al., 2017; Harrison et al., 2016; Sabuncu et al., 2012), finding alterations associated with increased genetic burden. Some of the studies have also used disease pathways to inform the PRS (Caspers et al., 2020; Ahmad et al., 2018). A few studies have also identified alterations in the brain of young AD-risk carriers. The hippocampal volume and the fractional anisotropy of the right cingulum are altered in young adults with increased risk of developing AD (Foley et al. 2017), and their precuneal volume is reduced (Li et al., 2018). Increased functional connectivity and hippocampal activation in a memory task was observed in the DMN of young, cognitively normal *APOE4* carriers (Filippini et al., 2009). Young *APOE4* carriers also showed increased activation (measured via fMRI) in the medial temporal lobe compared to non-carriers, while performing a memory task (Dennis et al., 2010).

Despite the evidence that a) there are alterations in the brain networks of AD patients, and b) there are functional and structural changes in the brains of young adults at risk of developing AD, the structural brain *networks* of young adults at risk of AD have not been studied. Our work fills that gap, by investigating structural brain networks of young adults at different risks of developing AD, where the risk is evaluated both via GWAS and via specific risk pathways. We hypothesise that the localized alterations in the structure of the brain of young adults at risk of AD would present themselves as changes in their structural brain networks. We investigate the network corresponding to the whole-brain connectome, as well as the DMN, the limbic and visual subnetworks, because those subnetworks are known to be affected in AD (Power et al., 2011; Deng et al. 2016; Hansson et al., 2017; Badhwar et al., 2017; Wang et al. 2019). We also investigate the hubs of the whole-brain connectome and their interconnectivity.

### 1.1 Hypotheses

We hypothesise that increased risk of AD would lead to increased characteristic path length for those networks, and an increased mean clustering coefficient, in agreement with the alterations these measures present in AD. We also hypothesize that the interconnectivity of the hubs would be reduced for increased risk of AD. Given the young age of the participants, we expect any observed alterations to be small. Any identified changes could be followed up in a longitudinal study of the same cohort, and possibly lead to important biomarkers that indicate disease onset or progression, or inform early preventative interventions in adults at risk of AD.

## 2. Materials and Methods

### 2.1 Participants

The Avon Longitudinal Study of Parents and Children (ALSPAC) is a pregnancy and birth cohort established to identify the factors influencing child health and developmental outcomes. Pregnant women resident in Avon, UK with expected dates of delivery 1st April 1991 to 31st December 1992 were invited to take part in the study. The initial number of pregnancies enrolled is 14,541 (for these at least one questionnaire has been returned or a “Children in Focus” clinic had been attended by 19/07/99). Of these initial pregnancies, there was a total of 14,676 foetuses, resulting in 14,062 live births and 13,988 children who were alive at 1 year of age.

Between the ages of 18 to 24 years, a subset of ALSPAC offspring were invited to participate in three different neuroimaging studies; the ALSPAC Testosterone study (Liao et al., 2021; Patel et al., 2020; n= 513, mean age at attendance 19.62 years, range 18.00 to 21.50 years), the ALSPAC Psychotic Experiences (PE) study (Fonville et al., 2015; Drakesmith et al., 2015; Drakesmith et al., 2016; Drakesmith et al., 2019; n=252, mean age at attendance 20.03 years, range 19.08 to 21.52 years), and the ALSPAC Schizophrenia Recall-by-Genotype (SCZ-RbG) study (Lancaster et al., 2019; n=196, mean age at attendance 22.75 years, range 21.12 to 24.55 years). Scanning protocols were harmonised across sub-studies where possible, and all data were acquired at Cardiff University Brain Research Imaging Centre (CUBRIC).

We analysed data from 562 individuals (mean age 19.81 years, SD 0.02 years; 62% male) from those ALSPAC neuroimaging studies (Boyd et al., 2013; Fraser et al., 2013; Sharp et al., 2020). Please note that the study website contains details of all the data that is available through a fully searchable data dictionary and variable search tool (http://www.bristol.ac.uk/alspac/researchers/our-data). Written informed consent was collected for all participants in line with the Declaration of Helsinki. Ethical approval for the neuroimaging studies was received from the ALSPAC Ethics and Law Committee and the local NHS Research Ethics Committees. Informed consent for the use of data collected via questionnaires and clinics was obtained from participants following the recommendations of the ALSPAC Ethics and Law Committee at the time.

### 2.2 MRI acquisition

MRI data were acquired using a GE HD× 3T system (GE Healthcare, Milwaukee W1) at CUBRIC. Axial T1-weighted images were acquired using a 3D fast spoiled gradient recalled sequence (TR = 8ms, TE = 3ms, TI = 450ms, flip angle = 20°, matrix size = 256 × 192 × 159) to aid co-registration. Diffusion-weighted images were acquired with a twice refocused spin-echo echo-planar imaging sequence parallel to the anterior-posterior commissure and the acquisition was peripherally gated to the cardiac cycle. Data were collected from 60 slices of 2.4 mm thickness (FOV=230 mm, matrix size 96 × 96, TE = 87 ms, b-values 0 and 1200 s/mm^2^) using parallel imaging (ASSET factor = 2) encoding along 30 isotopically distributed directions according to vectors taken from the International Consortium for Brain Mapping protocol (Jones et al., 1999). For 219 of those participants, the diffusion-weighted images were acquired using 60 directions. For those participants, a subsample of the optimal 30 directions were used, alongside the first three images with b-value equal to 0 (see Foley et al., 2018, for further details; Afzali et al., 2021; Jones et al., 1999).

### 2.3 Data processing and tractography

Data pre-processing was performed as described by Foley et al. (2018). To summarise, T1 structural data were down-sampled to 1.5 × 1.5 × 1.5 mm^3^ resolution. Eddy-current and participant motion correction were performed with an affine registration to the non-diffusion-weighted images (Leemans and Jones, 2009) with appropriate reorienting of the encoding vectors. Echo-planar imaging of the diffusion-weighted data was performed, warping the data to the down-sampled T1-weighted images (Irfanoglu et al., 2012). RESTORE (Chang et al., 2005), RESDORE (Parker et al., 2013a) and free water correction (Pasternak et al., 2009) algorithms were run. Whole-brain tractography was performed for each data set using the damped Richardson-Lucy pipeline (Dell’Acqua et al., 2010) which has been shown to produce a reliable tractogram in cases of crossing fibers, and in-house MATLAB code (Parker et al., 2013b). The criteria used for termination of the tracts were: angle threshold of >45°, fibre orientation density function peak < 0.05 and fractional anisotropy <0.2.

### 2.4 Network construction

We used the Automated Anatomical Labelling (AAL) (Tzourio-Mazoyer et al., 2002) to define the 90 cortical and subcortical areas of the cerebrum that correspond to the nodes of the structural networks. The WM tracts linking those areas are the connections, or edges, of the networks. The network generation was performed in ExploreDTI-4.8.6 (Leemans et al., 2009). We generated two connectivity matrices for each participant, one in which the edges are weighted by the number of streamlines (NS) and one in which they are weighted by the mean fractional anisotropy (FA) of the diffusion tensor along the streamlines of the tracts. Both these metrics have been shown to result in measures of connectivity that exhibit heritability (Arnatkeviciute et al., 2020), repeatability (Yuan et al., 2018; Roine et al., 2019; Messaritaki et al., 2019; Dimitriadis et al., 2021) and functional relevance (Honey et al., 2009; Goni et al., 2014; Messaritaki et al., 2021). To reduce the possible number of false connections, structural connections reconstructed with 5 or fewer streamlines were discarded from the analysis. Furthermore, to avoid our results being dependent on this choice of threshold, the analysis was repeated for this threshold being from 1 to 12 streamlines. A graphical representation of this part of the analysis is shown in Fig. 1.

**Figure 1:**
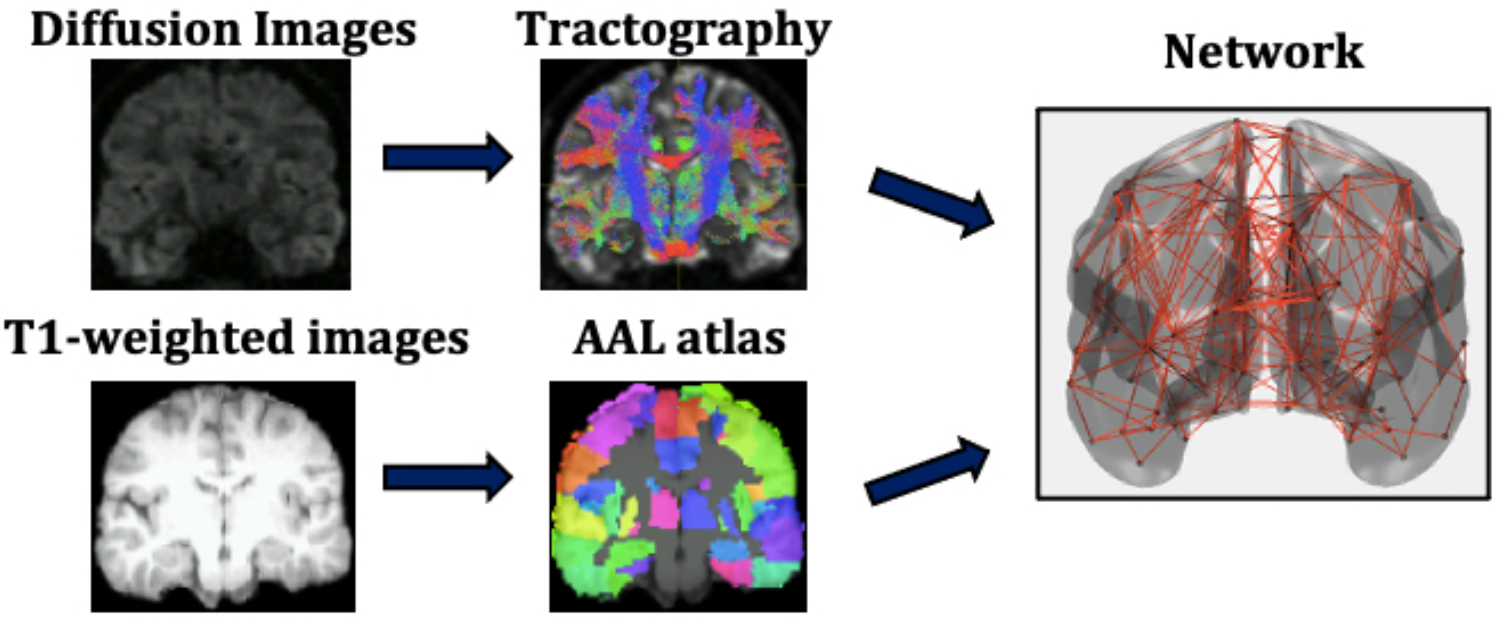
Analysis that leads from the MR images to the structural brain networks. This analysis is repeated for each participant individually.

In addition to the whole-brain connectome, we derived the DMN, the limbic subnetwork and the visual subnetwork, by selecting the edges that connect only the nodes in those subnetworks. The AAL atlas regions for the subnetworks are listed in Table 1 (Power et al., 2011).

**Table 1:**
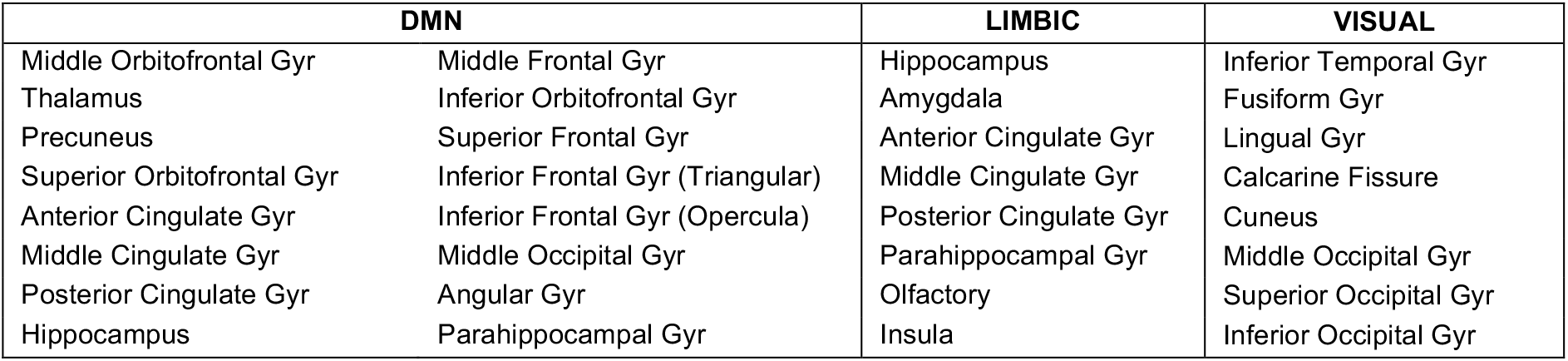
Nodes of the AAL atlas included in the DMN, limbic and visual subnetworks. The nodes from both the left and right hemispheres are included.

| DMN |  | LIMBIC | VISUAL |
| --- | --- | --- | --- |
| Middle Orbitofrontal Gyr | Middle Frontal Gyr | Hippocampus | Inferior Temporal Gyr |
| Thalamus | Inferior Orbitofrontal Gyr | Amygdala | Fusiform Gyr |
| Precuneus | Superior Frontal Gyr | Anterior Cingulate Gyr | Lingual Gyr |
| Superior Orbitofrontal Gyr | Inferior Frontal Gyr (Triangular) | Middle Cingulate Gyr | Calcarine Fissure |
| Anterior Cingulate Gyr | Inferior Frontal Gyr (Opercula) | Posterior Cingulate Gyr | Cuneus |
| Middle Cingulate Gyr | Middle Occipital Gyr | Parahippocampal Gyr | Middle Occipital Gyr |
| Posterior Cingulate Gyr | Angular Gyr | Olfactory | Superior Occipital Gyr |
| Hippocampus | Parahippocampal Gyr | Insula | Inferior Occipital Gyr |

### 2.5 Graph theory and network analysis

The Brain Connectivity Toolbox (BCT, Rubinov and Sporns, 2010) was used to calculate graph theoretical metrics for the structural brain networks of all participants. A detailed description of graph theoretical metrics is provided by Rubinov and Sporns, 2010, but we provide here a brief explanation of the ones we use, for completeness.

The *clustering coefficient* of a node is equal to the number of existing edges among the neighbours of the node divided by the number of all possible edges and is a measure of how interconnected the node’s neighbours are. The *degree* of a node is the number of edges that stem from that node. The *betweenness centrality* of a node is the number of shortest paths (connecting pairs of nodes) that the node belongs to in the network. The *nodal strength* is the sum of the weights of the edges stemming from a node. These four graph theoretical metrics are node-specific. To derive network-wide measures, their mean values over all the nodes in the network are used. The *characteristic path length* of a network is the mean value of the steps along the shortest paths that connect all possible pairs of nodes in the network. The *global efficiency* of the network is proportional to the sum of the inverse shortest path lengths over all pairs of nodes in the network and is related to how efficiently the nodes of the network can exchange information. In contrast to the previous measures mentioned, the characteristic path length and the global efficiency are network-wide, rather than node-specific, measures. Finally, the *local efficiency* of a node is calculated the same way as the global efficiency of the subnetwork that consists of the node’s neighbours.

**Figure 2:**
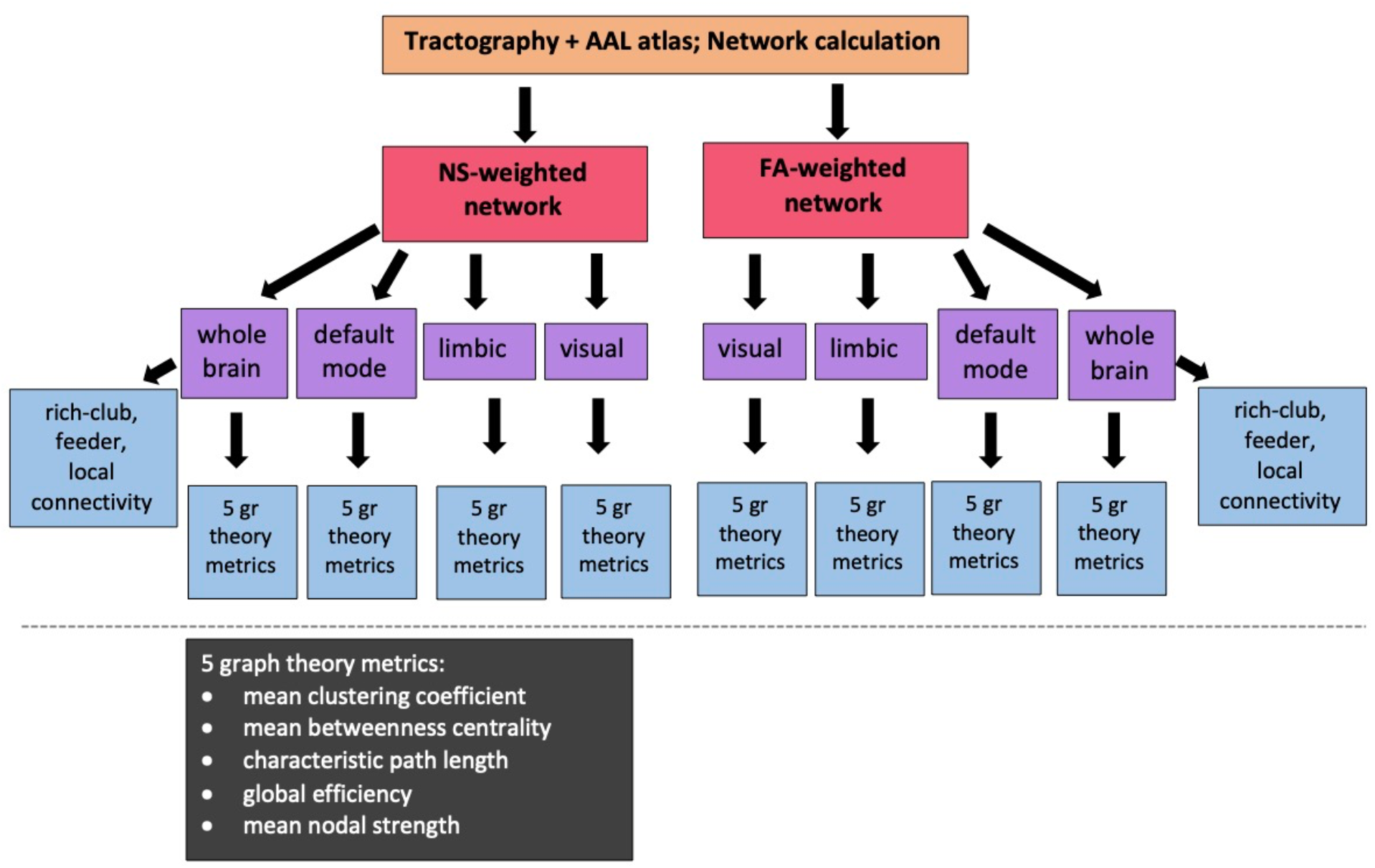
Diagram showing the sub/networks used in our analysis and the graph theoretical and connectivity metrics that are correlated with the PRS. NS = number of streamlines, FA = fractional anisotropy of the diffusion tensor.

For our analysis, we calculated the mean clustering coefficient, mean betweenness centrality, characteristic path length, global efficiency and mean nodal strength. The expectation is that, if changes to the topological organisation are a result of increased risk of developing AD, then the mean clustering coefficient, global efficiency and mean nodal strength will decrease, and the characteristic path length will increase, for increased risk. In order to remove metrics that represent redundant information from our analysis, we calculated the Pearson correlation between all pairs of graph theoretical metrics for each network and excluded from further analysis metrics that exhibited correlation coefficients of 0.85 or higher.

In order to investigate the hubs of the networks, we also calculated the local efficiency and the degree of each node. This allowed us to calculate the hub-score, or hubness, of each node for the whole-brain network. Instead of using a single measure for identifying hubs (for example only the node degree or only the betweenness centrality as is sometimes done), we used a composite measure as proposed by Betzel et al. (2014). Specifically, we normalized the node degree, nodal strength, betweenness centrality and local efficiency for each participant – this was in order for all four metrics to be equally weighted in the hubness calculation and was done by dividing the values of each metric across nodes by the largest value. We then averaged the normalized values for each node. That average was the *hubness* of the node.

The hubness of each node was averaged over all participants, to derive the mean node hubness. Hub nodes were defined as those with mean node hubness greater or equal to the average of the mean node hubnesses plus one standard deviation, according to van den Heuvel and Sporns (2011). The hub nodes comprise a *rich club* of nodes. The rich-club connectivity was calculated for each participant by summing the strength of the edges that connect the hub nodes only. The *feeder* connections, i.e., the connections that link one hub node and one nonhub node, were also identified. The feeder connectivity was also calculated for each participant, as the sum of the strength of the feeder connections. Finally, the *local* connections were identified as the connections that link non-hub nodes only. The local connectivity was the sum of the strength of the local connections. We stress that the rich-club, feeder and local connectivities are defined for the whole-brain network.

### 2.6 Polygenic risk score calculation

Genome data were provided by the University of Bristol. ALSPAC participants were genotyped using the Illumina HumanHap550 quad genome-wide SNP genotyping platform by 23andMe subcontracting the Wellcome Trust Sanger Institute (WTSI, Cambridge, UK) and the Laboratory Corporation of America (Burlington, North Carolina, USA). Participants were excluded from analysis if they had minimal or excessive heterozygosity, genotyping completeness < 97%, or if they were of non-European ethnicity. Quality control parameters were as follows: Minor allele frequency (MAF) > 0.01; Individual call rate > 95%, Hardy Weinberg Equilibrium (HWE) (P > 5×10^−7^). Polygenic risk scores were calculated according to the International Schizophrenia Consortium method (Purcell et al., 2009). Training data were taken from the latest genetic metaanalysis of Alzheimer’s disease (Kunkle et al, 2019) comprising of 94,437 cases and controls. In our sample, SNPs with low MAF < 0.1 and imputation quality <0.9 were removed. Data were then pruned for SNPs in linkage disequilibrium (LD) using genetic data analysis tool PLINK (Chang et al., 2015) using the clumping function (--clump). This aimed to remove SNPs in LD within a 500 kilobase window, retaining only the most significantly associated SNPs. Scores were generated in PLINK using the –score command. We note that *APOE* has a *p*-value of around 7×10^−44^ in most Alzheimer’s GWAS, and it explains almost as much variance in the phenotype as all the other loci combined. Therefore, *APOE4* carriers are invariably in the highest deciles of the polygenic score.

To compute pathway-specific PRS, nine pathway groups were taken from Kunkle et al. (2019), who matched lists of SNPs to genes and tested them for enrichment within gene functional categories. The pathway groups were as follows: protein-lipid complex assembly, regulation of beta-amyloid formation, protein-lipid complex, regulation of amyloid precursor protein catabolic process, tau protein binding, reverse cholesterol transport, protein-lipid complex subunit organisation, plasma lipoprotein particle assembly and activation of the immune response. The lists of SNPs were matched to SNPs in our target dataset. Then the data was clumped and scored as described above.

A previous study found that an AD PRS computed with *p*-value threshold (P_T_) of 0.001 explained the most variance in structural (non-network) neuroimaging phenotypes of healthy young adults (Foley et al. 2017). Therefore, our primary analysis used P_T_=0.001 to select relevant SNPs from the discovery sample. For our secondary analysis, 7 different progressive training P_T_S were computed (0.00001; 0.0001; 0.01; 0.05; 0.1; 0.3; and 0.5). Lower P_T_ indicates that SNPs are more significantly associated with AD case status in the training dataset. Two versions of each score were calculated, including and excluding the *APOE* locus. This was done to assess the effect of PRS without *APOE* and the effect of *APOE* within the PRS.

Through this method, we ended up with 20 different PRS: genome-wide with and without *APOE*,and each of the nine pathway-specific PRS with and without *APOE*. Each of these PRS further corresponds to 8 values for the P_T_S, as described above.

### 2.7 Statistical analyses

Correlations between graph theoretical metrics and the genome-wide PRS (*APOE* included) and the nine pathway-specific PRS (*APOE* included) were calculated in MATLAB (MATLAB and Statistics Toolbox Release, 2015b and 2021a; The MathWorks, Inc, Massachusetts, United States). Correlations were also calculated between the rich-club, feeder and local connectivity versus the 10 PRS scores. The participant gender and the diffusion scan type (30 vs 60 diffusion gradient directions) were controlled for by using partial correlations. Data points that had Cook’s distance higher than 3 times the mean Cook’s distance (Cook, 1977) were removed from the calculation. Our primary analysis used P_T_ = 0.001. Resulting *p*-values were corrected for multiple comparisons using false-discovery-rate (FDR) correction (Benjamini and Yekutieli, 2005). The correction was applied over the graph theoretical metrics of all four networks, the rich-club, feeder and local connectivities, and the 10 PRS (i.e., the genome-wide plus the 9 pathway-specific ones) for each P_T_. If a significant association was found between a PRS and the graph theoretical metrics or connectivities, correlations were also calculated with the PRS excluding the *APOE* locus, to assess whether the correlations were purely due to that locus. To exclude the possibility that our results are confounded by population stratification, we repeated our analyses using the first ten principal components derived from common alleles as covariates.

We also looked at the rest of the P_T_ thresholds, as is standard practice (de Leeuw et al., 2015; Purcell et al., 2009). In particular, the investigations for the higher values of P_T_ are justified, because it is likely that many loci that show only nominal association with disease status are actually involved in the pathological process. This was demonstrated by Escott-Price et al., (2015) who found that the highest prediction accuracy was given by a PRS which included SNPs from 0.5 and below (AUC = 78.2%, 95% confidence interval: 77 – 80 %). To control for multiple comparisons in this case, we calculated the permutation-corrected *p*-values via the *minP* procedure (Rempala and Yang, 2013; Westfall and Young, 1993).

## 3. Results

### 3.1 Networks

The whole-brain, default mode, limbic and visual subnetworks for one participant are shown in Fig. 3 (NS-weighted networks) and Fig. 4 (FA-weighted networks). The relative strength of the connections depends on the edge-weighting and has an impact on the graph theoretical metrics of the networks. Given the differences observed between NS- and FA-weighted networks, performing the analysis for both these edge-weightings is warranted.

**Figure 3:**
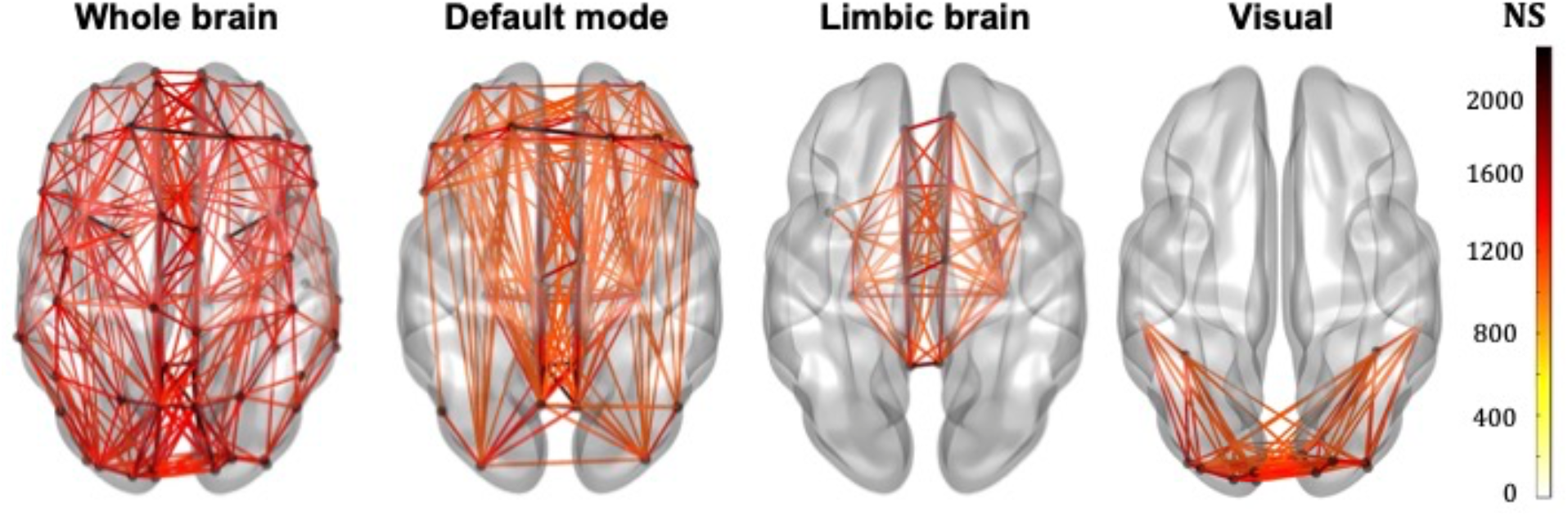
Whole-brain, DMN, limbic and visual subnetworks, for NS-weighted networks, from the data of one participant. The lines represent the edges (connections) between brain areas.

**Figure 4:**
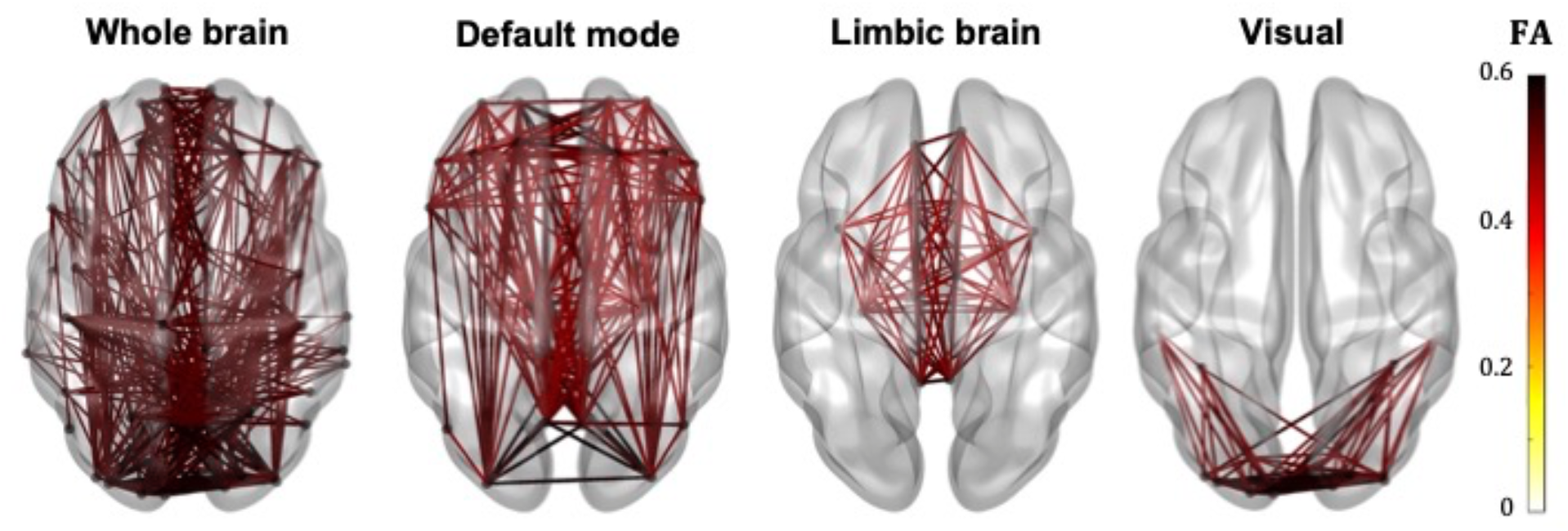
Whole-brain, DMN, limbic and visual subnetworks, for FA-weighted networks, from the data of one participant. The lines represent the edges (connections) between brain areas.

The correlation coefficients between graph theoretical metrics of the networks and the related *p*-values are given in Table 2a (for the NS-weighted networks) and 2b (for the FA-weighted networks). Based on these, we selected the metrics to be used in subsequent analysis, which are summarised in Table 3.

**Table 2a:**
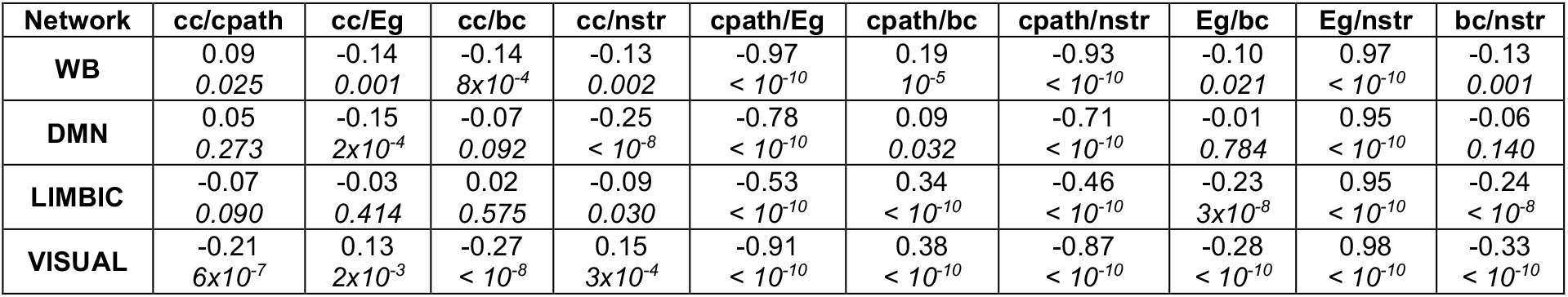
Correlation coefficients and *p*-values (the latter in italics) for the graph theoretical metrics of the NS-weighted networks. cc = mean clustering coefficient, cpath = characteristic path length, Eg = global efficiency, nstr = mean nodal strength, bc = mean betweenness centrality.

**Table 2b:** Correlation coefficients and *p*-values (the latter in italics) for the graph theoretical metrics of the NS-weighted networks. cc = mean clustering coefficient, cpath = characteristic path length, Eg = global efficiency, nstr = mean nodal strength, bc = mean betweenness centrality.

| Network | cc/cpath | cc/Eg | cc/bc | cc/nstr | cpath/Eg | cpath/bc | cpath/nstr | Eg/bc | Eg/nstr | bc/nstr |
| --- | --- | --- | --- | --- | --- | --- | --- | --- | --- | --- |
| WB | -0.35<br>*< 10<sup>-10</sup>* | 0.37<br>*< 10<sup>-10</sup>* | -0.31<br>*< 10<sup>-10</sup>* | 0.44<br>*< 10<sup>-10</sup>* | -0.99<br>*< 10<sup>-10</sup>* | 0.69<br>*< 10<sup>-10</sup>* | -0.91<br>*< 10<sup>-10</sup>* | -0.65<br>*< 10<sup>-10</sup>* | 0.91<br>*< 10<sup>-10</sup>* | -0.85<br>*< 10<sup>-10</sup>* |
| DMN | -0.44<br>*< 10<sup>-10</sup>* | 0.45<br>*< 10<sup>-10</sup>* | -0.45<br>*< 10<sup>-10</sup>* | 0.53<br>*< 10<sup>-10</sup>* | -0.98<br>*< 10<sup>-10</sup>* | 0.78<br>*< 10<sup>-10</sup>* | -0.92<br>*< 10<sup>-10</sup>* | -0.73<br>*< 10<sup>-10</sup>* | 0.93<br>*< 10<sup>-10</sup>* | -0.90<br>*< 10<sup>-10</sup>* |
| LIMBIC | -0.56<br>*< 10<sup>-10</sup>* | 0.55<br>*< 10<sup>-10</sup>* | -0.57<br>*< 10<sup>-10</sup>* | 0.63<br>*< 10<sup>-10</sup>* | -0.98<br>*< 10<sup>-10</sup>* | 0.81<br>*< 10<sup>-10</sup>* | -0.95<br>*< 10<sup>-10</sup>* | -0.76<br>*< 10<sup>-10</sup>* | 0.94<br>*< 10<sup>-10</sup>* | -0.92<br>*< 10<sup>-10</sup>* |
| VISUAL | -0.43<br>*< 10<sup>-10</sup>* | 0.40<br>*< 10<sup>-10</sup>* | -0.33<br>*< 10<sup>-10</sup>* | 0.41<br>*< 10<sup>-10</sup>* | -0.98<br>*< 10<sup>-10</sup>* | 0.78<br>*< 10<sup>-10</sup>* | -0.96<br>*< 10<sup>-10</sup>* | -0.70<br>*< 10<sup>-10</sup>* | 0.96<br>*< 10<sup>-10</sup>* | -0.83<br>*< 10<sup>-10</sup>* |

**Table 3:** Metrics used in the analysis for each network.

|  | **NS-weighted** | **FA-weighted** |
| --- | --- | --- |
| **Whole-brain** | mean clustering coefficient<br>mean betweenness centrality<br>mean nodal strength<br>rich-club connectivity<br>feeder connectivity<br>local connectivity | mean clustering coefficient<br>mean betweenness centrality<br>mean nodal strength<br>rich-club connectivity<br>feeder connectivity<br>local connectivity |
| **Default-mode** | mean clustering coefficient<br>mean betweenness centrality<br>mean nodal strength<br>characteristic path length | mean clustering coefficient<br>mean betweenness centrality<br>mean nodal strength |
| **Limbic** | mean clustering coefficient<br>mean betweenness centrality<br>mean nodal strength<br>characteristic path length | mean clustering coefficient<br>mean betweenness centrality |
| **Visual** | mean clustering coefficient<br>mean betweenness centrality<br>mean nodal strength | mean clustering coefficient<br>mean betweenness centrality<br>mean nodal strength |

### 3.2 Whole-brain connectome

No statistically significant correlations between graph theoretical metrics of the whole-brain network and the PRSs were found to survive multiple comparison correction.

### 3.3 Default-mode network

For our primary analysis (P_T_ = 0.001), no statistically significant correlations between the PRS and the graph theoretical metrics of the DMN survived multiple-comparison correction. The following correlations, however, did survive multiple comparison correction:

For P_T_ = 0.3, the mean nodal strength of the NS-weighted DMN was correlated with the genomewide PRS, including *APOE*, (r = −0.14, *p* = 1.5×10^−3^). When the *APOE* locus was excluded from the analysis, the correlation persisted (r = −0.14, *p* = 1.6×10^−3^). The correlations also persisted when the analysis was repeated for NS thresholds between 1 and 12.

For P_T_ = 0.01, the mean betweenness centrality of the FA-weighted DMN was correlated with the activation of the immune response PRS, including *APOE*, (r = −0.16, *p*=1.2×10^−4^). When the *APOE* locus was excluded from the analysis, the correlation persisted (r = −0.15, *p*=4.5×10^−4^). The correlations also persisted when the analysis was repeated for NS thresholds between 1 and 12.

Repeating the analyses for the DMN using the first ten principal components derived from common alleles as covariates did not change these results.

All these results are shown in Fig. 5.

**Figure 5:**
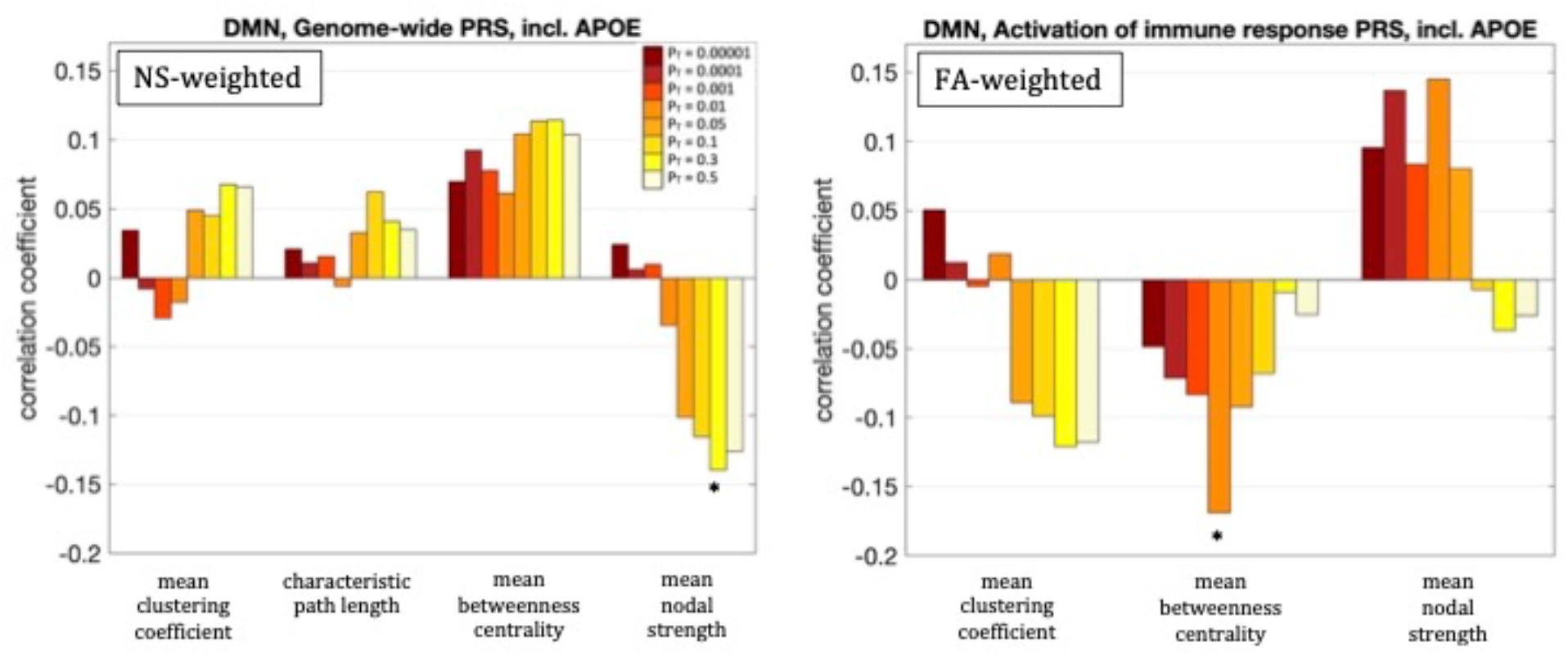
Correlation coefficients between the graph theoretical metrics of the default-mode network and the genomewide PRS including *APOE* for the 8 different values of P_T_. The asterisk indicates the instances in which the *p*-value survived multiple comparison correction.

### 3.4 Limbic subnetwork

No statistically significant correlations between graph theoretical metrics of the limbic subnetwork and the PRSs were found to survive multiple comparison correction.

### 3.5 Visual subnetwork

For our primary analysis (P_T_ = 0.001), no statistically significant correlations between the PRS and the graph theoretical metrics of the visual subnetwork survived multiple-comparison correction. The following correlations, however, did survive multiple comparison correction:

The mean nodal strength of the NS-weighted visual subnetwork was correlated with the genome-wide PRS, including *APOE*, for P_T_=0.1, 0.3 and 0.5. The correlation coefficients were r = −0.17, −0.18 and −0.19, for the 3 values of P_T_ respectively, while the *p*-values were 8.4×10^−5^, 4.1×10^−5^ and 1.3×10^−5^ respectively. When the analysis was repeated with the *APOE* locus excluded, the correlations persisted. Specifically, the correlation coefficients were: −0.15, −0.17 and −0.18, while the *p*-values were 7.6×10^−4^, 1.4×10^−4^, 2.9×10^−5^, for the 3 values of P_T_ respectively. The correlations also persisted when the analysis was repeated for NS thresholds between 1 and 12.

The mean clustering coefficient of the NS-weighted visual subnetwork was correlated with the tau protein binding PRS, including *APOE*, for P_T_=0.3 and 0.5. The correlation coefficients were r = −0.14, while the *p*-values were 1.4×10^−3^ for both P_T_S. When the analysis was repeated with the *APOE* locus excluded, the significance of the correlations disappeared, with the correlation coefficients being −0.02 and the *p*-values being 0.71. The correlations persisted, however, when the analysis was repeated for NS thresholds between 1 and 12.

The mean betweenness centrality of the NS-weighted visual subnetwork was correlated with the plasma lipoprotein particle assembly PRS, including *APOE*, for P_T_=0.3 and 0.5. The correlation coefficients were r = 0.15 and 0.16, for the 2 values of P_T_ respectively, while the *p*-values were 9.2×10^−4^ and 3.6×10^−4^ respectively. When the analysis was repeated with the *APOE* locus excluded, the correlations persisted, with the correlation coefficients being r = 0.12 and 0.13 for the two values of P_T_ respectively, and the *p*-values being 7.5×10^−3^ and 2.2×10^−3^ respectively. The correlations also persisted when the analysis was repeated for NS thresholds between 1 and 12.

Repeating the analyses for the DMN using the first ten principal components derived from common alleles as covariates did not change these results.

All these results are shown in Fig. 6.

**Figure 6:**
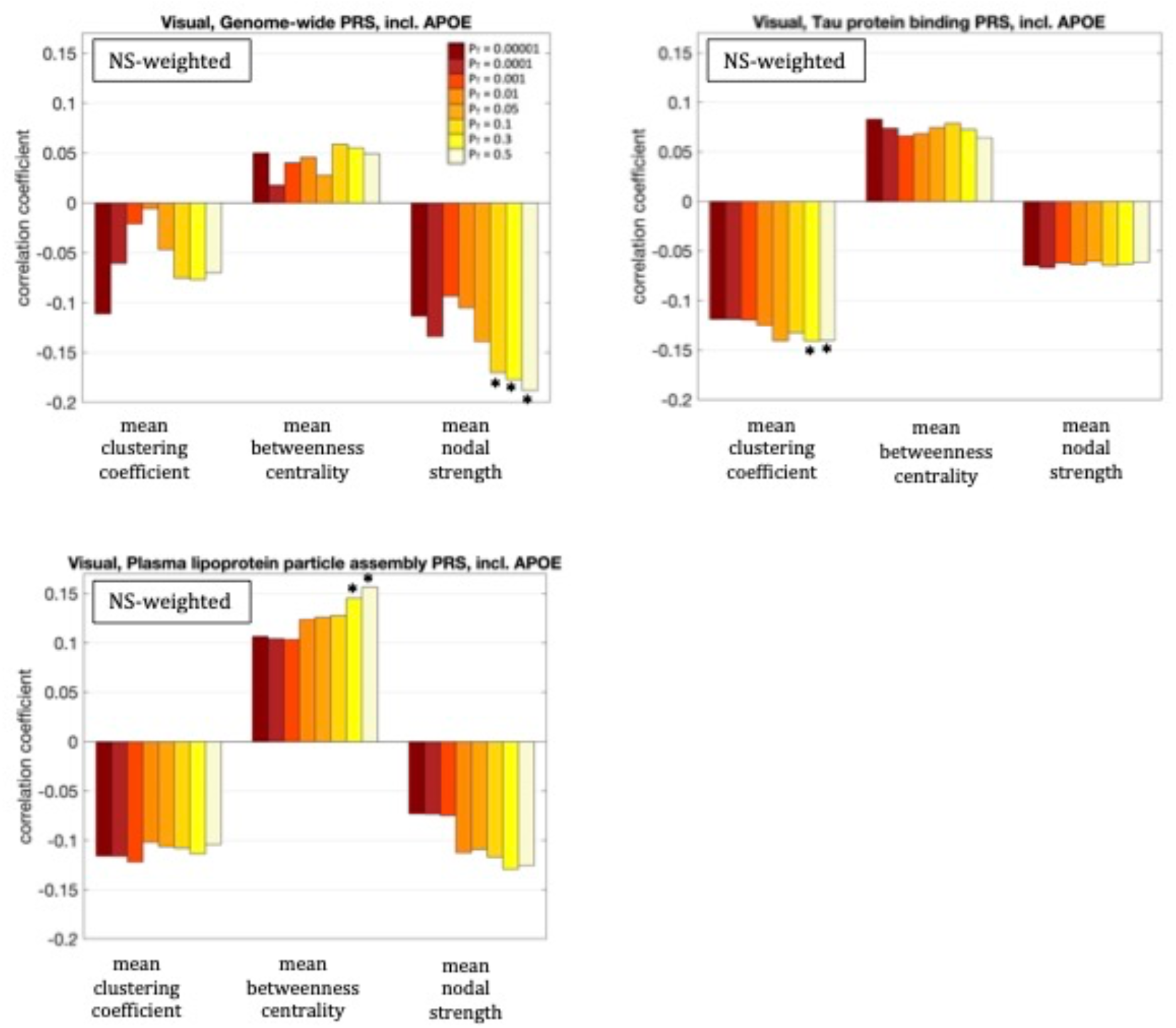
Correlation coefficients between the graph theoretical metrics of the visual subnetwork and the three PRSs for which those survived multiple comparison correction, for the 8 different values of P_T_. The asterisk indicates the instances in which the *p*-value survived multiple comparison correction.

### 3.6 Rich-club, feeder and local connectivity of the whole-brain network

Fig. 7 shows the nodes that are hubs for the NS-weighted and the FA-weighted networks. For the NS-weighted networks, the hubs were the left and right putamen, left and right precuneus, left and right hippocampus, left and right superior frontal gyrus, left middle occipital gyrus, left and right superior occipital gyrus, right calcarine sulcus and right caudate. For the FA-weighted networks, the hubs were the left and right putamen, left and right precuneus, left and right hippocampus, left and right superior frontal gyrus, left middle occipital gyrus, left calcarine sulcus, right superior parietal gyrus, left superior orbitofrontal gyrus and left superior occipital gyrus. We note that ten out of the 13 hubs were the same in the NS- and FA-weighted networks, while three differed.

**Figure 7:**
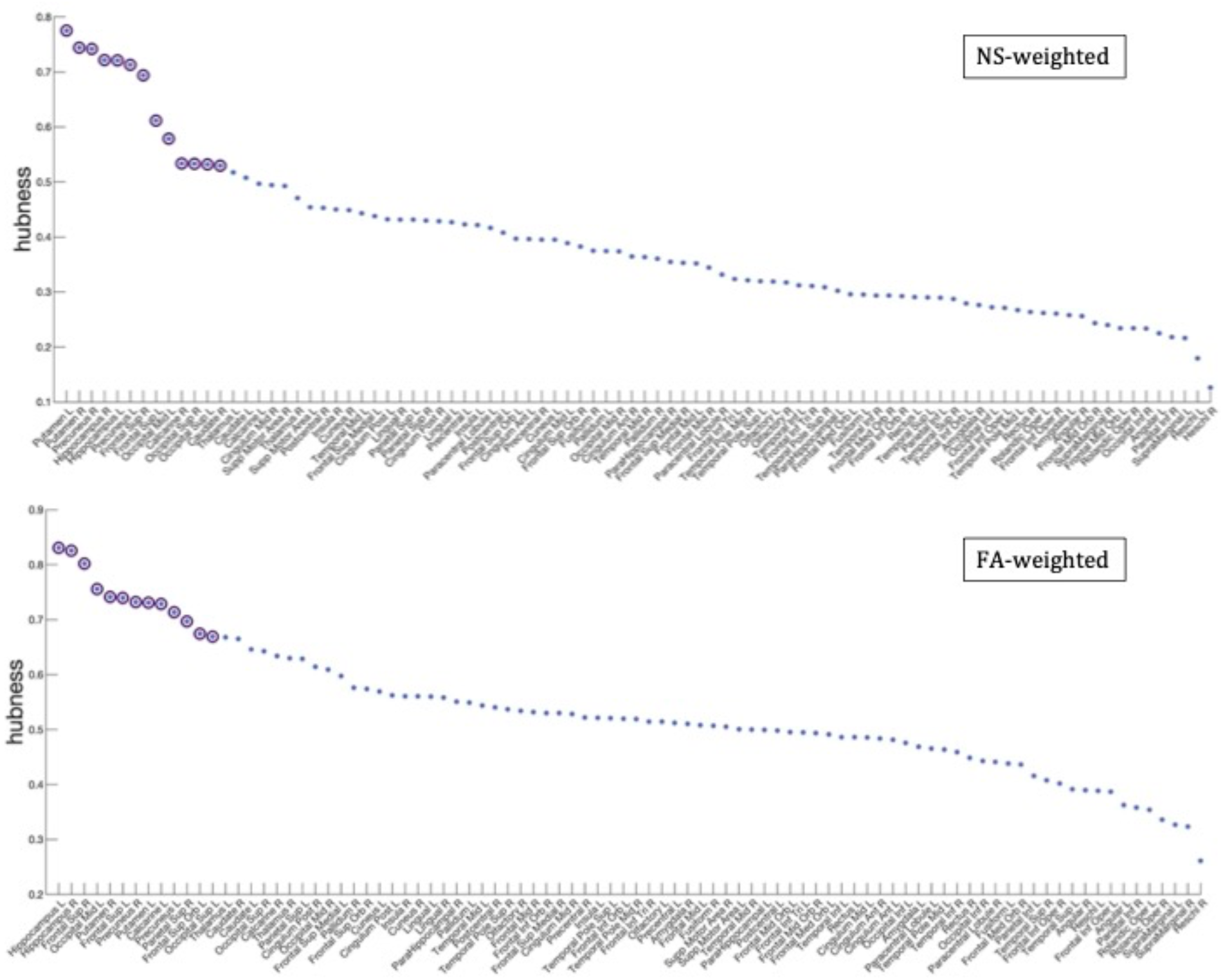
Hubness scores for the network nodes for the NS-weighted (top) and FA-weighted (bottom) networks. The purple circles indicate nodes that are hubs for the respective networks.

For our primary analysis (P_T_ = 0.001), no statistically significant correlations between the PRS and the rich-club, feeder or local connectivities of the whole-brain network survived multiple-comparison correction. The following correlations, however, did survive multiple comparison correction:

The rich-club connectivity of the NS-weighted whole-brain connectome was correlated with the genome-wide PRS, including *APOE*, for P_T_=0.3 and 0.5. The correlation coefficients were r = −0.16 and −0.15 for the two P_T_S respectively, while the p-values were 3.7×10^−4^ and 1.1×10^−3^ respectively. When the analysis was repeated with the *APOE* locus excluded, the correlations persisted, with the correlation coefficients being r = −0.15 and −0.14 for the two P_T_S respectively, and the *p*-values being 6×10^−4^ and 1.7×10^−3^ respectively. The correlations also persisted when the analysis was repeated for NS thresholds of 1 to 12.

The feeder connectivity of the NS-weighted whole-brain connectome was correlated with the genome-wide PRS, including *APOE*, for P_T_=0.3 and 0.5. The correlation coefficients were r = −0.14 and −0.15 for the two P_T_S respectively, while the p-values were 1.3×10^−3^ and 8.8×10^−4^ respectively. When the analysis was repeated with the *APOE* locus excluded, the correlations persisted, with the correlation coefficients being r = −0.14 and −0.13 for the two P_T_S respectively, and the *p*-values being 1.4×10^−3^ and 2.3×10^−3^ respectively. The correlations also persisted when the analysis was repeated for NS thresholds of 1 to 12.

All these results are shown in Fig. 8.

**Figure 8:**
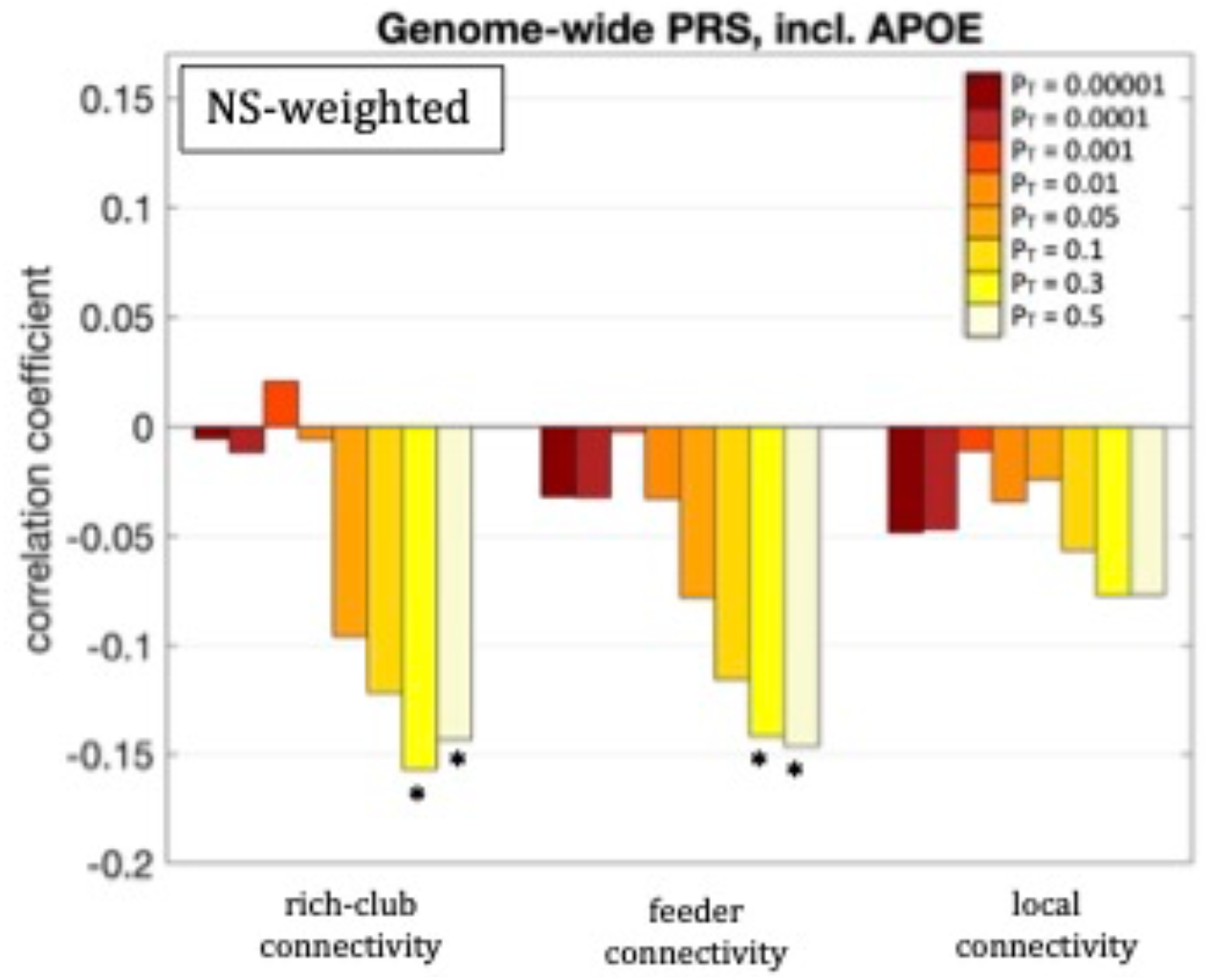
Correlation coefficients between the rich-club, feeder and local connectivities and the genome-wide PRS including *APOE*, for the 8 different values of P_T_. Asterisks indicate *p*-values that survived multiple comparison correction.

As mentioned earlier, the *minP* procedure was used to calculate the permutation-corrected *p*-values for the P_T_ thresholds. The exact *p*-values are given in Section A of the Supplementary Material. They remained statistically significant for all the cases above, with the exception of the correlation between the mean betweenness centrality of the visual network and the PRS for plasma lipoprotein particle assembly excluding APOE for P_T_ = 0.3, and the correlation between the feeder connectivity and the genome-wide PRS excluding APOE for P_T_ = 0.3. Additionally, the values of the correlation coefficients for the thresholds between 1 and 12 are given in Section B of the Supplementary Material.

## 4. Discussion

To the best of our knowledge, this is the first study to examine the relationship between AD PRS and network-based measures for the whole-brain structural connectome and subnetworks. We used a cohort of young participants to assess any potential early changes in the structural connectome. From a clinical perspective, using pathway-specific polygenic risk scores in addition to genome-wide ones is important, because it can pave the way for more targeted interventions based on the predicted pathway involvement and potentially allow clinical trials to stratify patients using their specific risk profiles.

Compared to the FA-weighted networks, using NS-weighted networks resulted in more statistically significant relationships between the PRS and structural network metrics, such as the graph theoretical metrics we employed and the connectivity strength between the rich-club, and feeder connections. Even though both the NS and the FA are routinely used to assign significance to the edges of structural networks, it has been argued (Huang and Ding, 2016) and proven experimentally (Messaritaki et al., 2021) that the NS is more relevant from a functional perspective to the network organization of the human brain compared to the FA. This may be contributing to the increased sensitivity of the NS in the differences observed in our study. Other metrics have also been used as edge-weights, such as the inverse radial diffusivity (Caeyenberghs et al., 2016; Messaritaki et al., 2022), which captures myelination and axonal packing and is, therefore, also meaningful is assessing connectivity. From a methodological point of view, this demonstrates that the selection of the metric for the edge weights can impact the results and, if not optimal, it can fail to reveal certain statistically significant relationships. As the capabilities of MRI to measure microstructural metrics evolves (Wolff et al., 1989; Mackay et al., 1994; Assaf and Basser, 2005; Zhang et al., 2012; Barazany et al., 2009), using these and other measures (e.g., myelin, axonal density and axon diameter) as edge weights should also be explored.

Our analysis identified statistically significant (after correction for multiple comparisons) correlations between graph theoretical metrics and PRS, present in the DMN. The negative correlation between the mean nodal strength and the genome-wide PRS for the NS-weighted DMN indicates that high genome-wide risk of AD results in lower nodal strength in that network. Furthermore, the fact that the correlation persisted when the *APOE* locus was removed from the analysis indicates that this relationship is a result of multiple genetic factors and not exclusively due to the *APOE* gene.

Our analysis also revealed statistically significant (after multiple-comparison correction) correlations between the graph theoretical metrics of the NS-weighted visual subnetwork and the PRSs. The negative correlation between the mean nodal strength and the genome-wide PRS (including *APOE*) implies weaker connectivity in the visual subnetwork of participants at higher risk of developing AD. The negative correlation between the mean clustering coefficient and the tau protein binding PRS (including *APOE*) indicates that participants at higher risk of developing AD through this pathway have less clustered communities in the visual subnetwork. The positive correlation between the mean betweenness centrality and the PRS for plasma lipoprotein particle assembly (including *APOE*) implies that, in participants at higher risk of developing AD, each node participates in more shortest paths and therefore the organisation of the visual subnetwork is less central compared to participants at low risk. The fact that the first and third of these correlations persisted when the *APOE* locus was excluded from the genetic risk calculation indicates that they are a result of multiple genetic factors, and not exclusively due to the *APOE* gene. The second correlation, however, appears to be driven predominantly by the *APOE* gene.

Studies of young adults with genetic predisposition to AD are still limited, and predominantly involve brain function rather than structure. As mentioned earlier, increased functional connectivity and hippocampal activation in a memory task was observed in the DMN of young, cognitively normal *APOE4* carriers (Filippini et al., 2009). This finding was not, however, replicated in a study by Mentink et al. (2021), which instead found that compared to non-carriers, *APOE4* young carriers had increased functional activation in facial-recognition areas during the encoding of subsequently recollected items. Young *APOE4* carriers also showed increased activation (measured via fMRI) in the medial temporal lobe compared to non-carriers, while performing a memory task (Dennis et al., 2010). In contrast to these studies which focused on the DMN, the majority of our findings pertained to the visual network. Additionally, observing small changes in the NS-weighted and FA-weighted structural networks does not necessarily imply the presence of measurable functional deficiencies (which also depends on the sensitivity of those functional studies). Functional connectivity is believed to be also reliant on a number of other microstructural metrics (such as myelination and axonal density) which could be compensating for changes present in the number of streamlines.

As mentioned earlier, alterations in the visual subnetwork of AD patients have been recently reported in the literature. For example, Deng et al. (2016) observed increased characteristic path length and clustering coefficient in the visual subnetwork (measured with BOLD fMRI) of AD patients. Badhwar et al. (2017) also observed decreased connectivity in the primary visual cortex of AD patients. Wang et al. (2019) observed impairments in the visual subnetwork of AD patients, as well as in patients with subjective cognitive decline, which is considered a prodromal stage of AD. This last result further supports the idea that alterations in the visual subnetwork can appear many years before AD diagnosis.

We also observed statistically significant correlations (after multiple-comparison correction) between the rich-club and feeder connectivities of the NS-weighted whole-brain network and the genome-wide PRS, including *APOE*. These negative correlations indicate that structural connections that involve at least one hub node are weaker in the brains of young participants at risk of developing AD. The relationships held when the *APOE* locus was excluded from the analysis, which indicates that the effect comes from genetic influences above and beyond *APOE*.

A few studies have reported altered connectivity of the rich-club and feeder edges in the structural brain networks of participants with Alzheimer’s disease and with MCI. Xue et al. (2020) recently observed reduced rich-club connectivity in patients with amnestic MCI compared to healthy age-matched controls, and reduced feeder and local connectivity in patients with amnestic MCI compared to participants with subjective cognitive decline. Cai et al. (2019) reported decreased feeder (and local) connection strength in the structural networks of AD patients compared to healthy controls. Our results are in line with these alterations in connectivity strength observed in AD and MCI patients.

It is interesting that we observed decreased clustering coefficient with increased AD risk, while AD studies (Deng et al., 2016; He et al., 2008; Lo et al., 2010) observed increased clustering coefficient in AD patients. However, it is not uncommon that a pattern of structural or functional metrics is observed in at-risk populations, for that pattern to be reversed when the pathology is realised. Specifically for AD for example, Koelewijn et al. (2019) observed that young *APOE* carriers exhibited hyperconnectivity in brain areas that were found, in the same work, to show hypoconnectivity in AD patients. We also note that the structural networks in He et al. (2008) were structural covariance networks rather than tractography-derived networks, and that Lo et al. (2010) had a small sample of 25 patients and 30 controls in their tractography study. Also, Deng et al. (2016) used functional MRI rather than diffusion MRI and had a much smaller sample than ours.

The rest of the graph theoretical metrics we investigated showed no statistically significant correlations after multiple comparison correction was applied. Recently, Foley et al. (2017) showed that there is a reduction in the fractional anisotropy of the right cingulum and a decrease in the left hippocampal volume of young adults at genetic risk of developing AD. In this context, our results imply that those alterations do not translate into changes in the structural brain networks and subnetworks of those young adults. We note, however, that the participants in that study included participants that were a few years older compared to those in our study.

The correlations observed in our analysis are small, in the range of 0.14 to 0.19 (in absolute value). This is to be expected, given that the cohort of our study consisted of young adults with normal brain function.

We note that the summary statistics used in PRS analysis were taken from a large discovery sample reported in the latest GWAS meta-analysis (Kunkle et al., 2019). Therefore, our risk estimates for AD loci are the best available. We computed PRS in our sample manually in PLINK, rather than using automated PRS tools. This gives us the ability to specify several exact parameters which can be difficult with automated PRS tools. Furthermore, these automatic tools have precalculated SNPs linkage disequilibrium scores. They often use only 1 million SNPs, whereas the current GWASes have 4-8 million SNPs available. Our study employed a relatively large sample size comprising participants of the same age (19.81 ±0.02 years old), therefore avoiding the confound of brain changes that are age related, and which are known to exist in young adults up to the age of at least 25 years. Furthermore, our study is the first to use disease pathway PRS to explore associations between biological pathways and underlying differences in structural brain connectivity. We used two different edge weights, NS and FA, in our analysis. These metrics are the most widely used in the literature to weigh the edges of structural networks. It is worth pointing out that some graph theoretical metrics are dependent on the choice of edge weighting (such as the clustering coefficient and the nodal strength), while others are not (such as the node degree). Furthermore, we could have calculated correlations for other graph theoretical metrics, however we chose to limit our choice as described in the Methods section, in order to avoid multiple comparison corrections forcing us to reject results that are truly statistically significant. We also note that changes in the topological properties of brain networks can be complex and due to a number of factors, such as, for example, volumetric changes, which could exhibit themselves as altered connectivity. Regarding the pathway-specific PRS, the accuracy of our results is limited by the current knowledge of pathway variants. Additionally, our study involved a geographically limited sample in which men are slightly overrepresented. Therefore, our results may not be representative of the general population. We note that the AAL atlas that we used is one of a few atlases that could have been used to conduct the analysis. Recent studies have shown how results from tractography studies could be dependent on the choice of atlas (Parker et al., 2014). Finally, participants of non-European ethnicity were excluded from the analysis because polygenic score analyses in populations with high genetic admixture are not valid. Even small differences in population genetics may lead to distinctive linkage disequilibrium (LD) structure and allele frequencies (Moskvina et al., 2010). Pruning, an essential part of PRS calculation, relies on LD structure to retain SNPs that are most associated with a trait while removing others that are closely linked. Where LD structure diverges, alternative SNPs will be selected. This means that ethnicity admixture must be avoided and comparisons between population groups using PRS are not valid. This further implies that the findings of our study may not generalize to other ethnicities. Around 2% of ALSPAC participants were non-white (Fraser et al, 2013).

## 5. Conclusion

Our results demonstrate that genetic burden is linked to changes in structural brain networks, both for the whole-brain connectome and the visual subnetwork, in young adults. The genome-wide PRS including *APOE* was linked to a reduction in the mean nodal strength of the visual subnetwork and of the rich-club and feeder connections of the whole-brain network. The plasma-lipoprotein particle assembly PRS including *APOE* was linked to an increase in the betweenness centrality of the visual subnetwork. Importantly, these relationships were still present, albeit slightly weaker, when the *APOE* locus was excluded from the analysis. This indicates that the search for AD biomarkers can benefit from the consideration of genetic risk above and beyond *APOE*. Different biomarkers could point to different pathway involvement, which could allow clinical trials to stratify patients accordingly. Specifically for the pathway-specific PRS, it is not currently known exactly how these biological processes relate to brain networks, and this is incredibly complicated to decipher. As such, this work points to possible directions that researchers can look into in future studies.

## Supplementary Material

### A. Permutation Tests

Below we give the *p*-values that resulted from the permutation testing, as described in Section 2.7. We report on the cases in which the correlations between graph theoretical metrics and PRS were statistically significant.

Permutation-corrected *p*-values for the correlation between mean nodal strength of the NS-weighted DMN and genome-wide PRS incl. APOE:

P_T_=0.3: 0.0438

Permutation-corrected *p*-values for the correlation between mean nodal strength of the NS-weighted DMN and genome-wide PRS excl. APOE:

P_T_=0.3: 0.0471

Permutation-corrected *p*-values for the correlation between mean betweenness centrality of the FA-weighted DMN and activation of the immune response PRS incl. APOE:

P_T_=0.01: 0.0083

Permutation-corrected *p*-values for the correlation between mean betweenness centrality of the FA-weighted DMN and activation of the immune response PRS excl. APOE:

P_T_=0.01: 0.0209

Permutation-corrected *p*-values for the correlation between mean nodal strength of the NS-weighted visual network and genome-wide PRS incl. APOE:

P_T_=0.1: 0.0062; P_T_=0.3: 0.0036; P_T_=0.5: 0.0014

Permutation-corrected *p*-values for the correlation between mean nodal strength of the NS-weighted visual network and genome-wide PRS excl. APOE:

P_T_=0.1: 0.0271; P_T_=0.3: 0.0093; P_T_=0.5: 0.0031

Permutation-corrected *p*-values for the correlation between mean clustering coefficient of the NS-weighted visual network and tau protein binding PRS incl. APOE:

P_T_=0.3: 0.0155; P_T_=0.5: 0.0157

Permutation-corrected *p*-values for the correlation between the mean betweenness centrality of the NS-weighted visual network and the plasma lipoprotein particle assembly PRS incl. APOE:

P_T_=0.3: 0.0102; P_T_=0.5: 0.0055

Permutation-corrected *p*-values for the correlation between the mean betweenness centrality of the NS-weighted visual network and the plasma lipoprotein particle assembly PRS excl. APOE:

P_T_=0.3: 0.0772; P_T_=0.5: 0.0333

Permutation-corrected *p*-values for the correlation between the rich-club connectivity of the NS-weighted whole-brain network and the genome-wide PRS incl. APOE:

P_T_=0.3: 0.0184; P_T_=0.5: 0.0346

Permutation-corrected *p*-values for the correlation between the rich-club connectivity of the NS-weighted whole-brain network and the genome-wide PRS excl. APOE:

P_T_=0.3: 0.0250; P_T_=0.5: 0.0492

Permutation-corrected *p*-values for the correlation between the feeder connectivity of the NS-weighted whole-brain network and the genome-wide PRS incl. APOE:

P_T_=0.3: 0.0291; P_T_=0.5: 0.0365

Permutation-corrected *p*-values for the correlation between the feeder connectivity of the NS-weighted whole-brain network and the genome-wide PRS excl. APOE:

P_T_=0.3: 0.0637; P_T_=0.5: 0.0463

### B. Correlation values for the different thresholds on the number of streamlines

The tables below give the correlation coefficients and the *p*-values for the different values of NS_thr_, where NS_thr_ is the maximum value of NS for tracts excluded from the brain network. For the case of the correlations not listed, the differences between the correlation coefficients / *p*-values for the 12 NS_thr_ were on the third significant digit, and for that reason we do not list them here.

Correlation between the mean betweenness centrality of the DMN and the activation of the immune response PRS (incl. APOE) for the FA-weighted networks, for P_T_=0.01:

| $NS_{thr}$ | 1 | 2 | 3 | 4 | 5 | 6 | 7 | 8 | 9 | 10 | 11 | 12 |
| --- | --- | --- | --- | --- | --- | --- | --- | --- | --- | --- | --- | --- |
| **Correlation coefficient** | -0.16 | -0.17 | -0.17 | -0.18 | -0.16 | -0.16 | -0.17 | -0.18 | -0.17 | -0.16 | -0.17 | -0.16 |
| **p-value** | $3 \times 10^{-4}$ | $10^{-4}$ | $9 \times 10^{-5}$ | $5 \times 10^{-5}$ | $1 \times 10^{-4}$ | $2 \times 10^{-4}$ | $2 \times 10^{-4}$ | $7 \times 10^{-5}$ | $2 \times 10^{-4}$ | $3 \times 10^{-4}$ | $2 \times 10^{-4}$ | $3 \times 10^{-4}$ |

Correlation between the mean clustering coefficient of the visual network and the tau protein binding PRS (incl. APOE) for the NS-weighted networks, for P_T_=0.3 and 0.5:

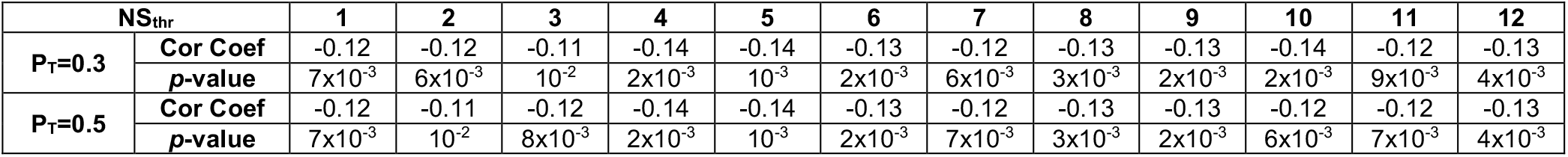

| NS<sub>thr</sub> |  | 1 | 2 | 3 | 4 | 5 | 6 | 7 | 8 | 9 | 10 | 11 | 12 |
| --- | --- | --- | --- | --- | --- | --- | --- | --- | --- | --- | --- | --- | --- |
| P<sub>T</sub>=0.3 | Cor Coef | -0.12 | -0.12 | -0.11 | -0.14 | -0.14 | -0.13 | -0.12 | -0.13 | -0.13 | -0.14 | -0.12 | -0.13 |
|  | p-value | 7×10<sup>-3</sup> | 6×10<sup>-3</sup> | 10<sup>-2</sup> | 2×10<sup>-3</sup> | 10<sup>-3</sup> | 2×10<sup>-3</sup> | 6×10<sup>-3</sup> | 3×10<sup>-3</sup> | 2×10<sup>-3</sup> | 2×10<sup>-3</sup> | 9×10<sup>-3</sup> | 4×10<sup>-3</sup> |
| P<sub>T</sub>=0.5 | Cor Coef | -0.12 | -0.11 | -0.12 | -0.14 | -0.14 | -0.13 | -0.12 | -0.13 | -0.13 | -0.12 | -0.12 | -0.13 |
|  | p-value | 7×10<sup>-3</sup> | 10<sup>-2</sup> | 8×10<sup>-3</sup> | 2×10<sup>-3</sup> | 10<sup>-3</sup> | 2×10<sup>-3</sup> | 7×10<sup>-3</sup> | 3×10<sup>-3</sup> | 2×10<sup>-3</sup> | 6×10<sup>-3</sup> | 7×10<sup>-3</sup> | 4×10<sup>-3</sup> |

Correlation between the rich club connectivity of the whole-brain network and the genome-wide PRS (incl. APOE) for the NS-weighted networks, for P_T_=0.3 and 0.5:

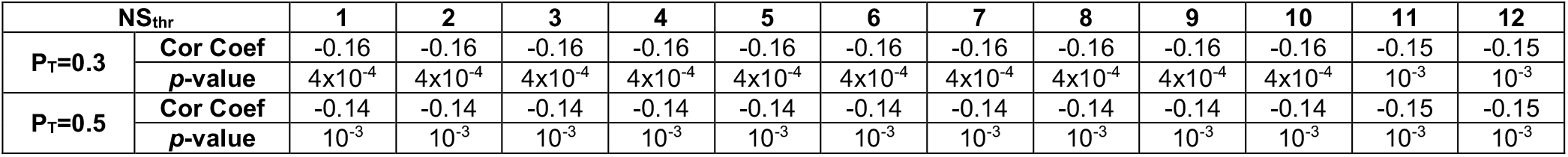

Correlation between the feeder connectivity of the whole-brain network and the genome-wide PRS (incl. APOE) for the NS-weighted networks, for P_T_=0.3 and 0.5:

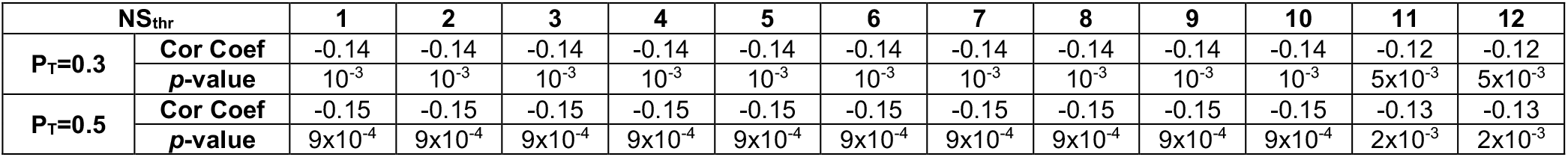

**Note**: EM and JRH are both last authors of the article.

## Acknowledgements

We are extremely grateful to the families that took part in this study, the midwives for their help in recruiting them, and the whole ALSPAC team, which includes interviewers, computer and laboratory technicians, clerical workers, research scientists, volunteers, managers, receptionists and nurses. We also wish to thank Dr. Maryam Afzali for useful discussions.

